# Drift-diffusion explains response variability and capacity for tracking objects

**DOI:** 10.1101/412650

**Authors:** Asieh Daneshi, Hamed Azarnoush, Farzad Towhidkhah, Ali Ghazizadeh

**Affiliations:** Biomedical Engineering Department, Amirkabir University of Technology (Tehran Polytechnic); Electrical Engineering Department, Sharif University of Technology; Center for Biological Intelligence, Sharif University of Technology

## Abstract

Being able to track objects that surround us is key for planning actions in dynamic environments. However, rigorous cognitive models for tracking of one or more objects are currently lacking. In this study, we asked human subjects to judge the time to contact (TTC) a finish line for one or two objects that became invisible shortly after moving. We showed that the pattern of subject responses had an error variance best explained by an inverse Gaussian distribution and consistent with the output of a biased drift-diffusion model. Furthermore, we demonstrated that the pattern of errors made when tracking two objects showed a level of dependence that was consistent with subjects using a single decision variable for reporting the TTC for two objects. This finding reveals a serious limitation in the capacity for tracking multiple objects resulting in error propagation between objects. Apart from explaining our own data, our approach helps interpret previous findings such as asymmetric interference when tracking multiple objects.

## 2. Introduction

Time to contact (TTC), which is the time it takes for an object to reach an observer or a particular place, is an important factor in a variety of real-world situations, such as catching and hitting balls in games, driving vehicles, or passing through a busy street. The ability to estimate TTC for one object has been assessed in several studies (for example see [1-14]).

The accuracy and precision of TTC estimation is related to the time perception ability, which is considered in several studies [6, 11, 15-24]. A type of task that is often considered in laboratory studies for TTC estimation is coincidence anticipation (CA) [5]. In CA tasks, subjects must make a simple response (e.g. press a button) when the moving object reaches a particular place, called contact point [5]. In an important type of CA tasks, often referred to as prediction motion (PM) tasks, the moving object disappears before reaching the contact point or hides behind an occluder. Then, after a specific time (here referred to as the extrapolation time), subject should indicate (often by pressing a button) the presumed time for the moving object to reach the contact point. The PM paradigm is used as a straightforward method to assess an individual’s ability to estimate absolute TTC (e.g., [4]). The main purpose of PM tasks is to understand how sensory and cognitive information are used to estimate TTC. To this aim, variables related to object’s motion, e.g., velocity, extrapolation distance and/or duration, are manipulated.

A few studies have looked at tracking multiple objects simultaneously. TTC judgment for two or more objects is required in a large number of everyday activities, such as crossing a multi-lane street or walking in crowded areas. One of the first studies which has considered multiple objects in TTC estimation tasks is done by Novak [25]. She presented multiple objects approaching a finish line, but the participants were asked to judge the TTC for only one of them. Most other studies presenting several objects simultaneously, have used relative judgment (RJ) tasks [3, 5, 26, 27]. In such tasks, subjects determine which of the two (or more) moving balls arrives first at a designated goal, after disappearing [5]. Here again, participants may compute and compare several TTCs, but they are asked for only one TTC estimation. The most important difference between RJ tasks and multi-object PM tasks is that for the former, participants may misestimate TTC for some or all objects, but still give the correct answer, as long as they preserve the perceived order of arrivals. In contrast, in multi-object PM tasks, the absolute accuracy of TTC judgment is assessed. Recently, Baures and colleagues have conducted some studies on the simultaneous estimation of the TTC for multiple objects [28-31]. However, their main focus has been on understanding how humans use their limited sources of attention to estimate several TTCs simultaneously, without considering the mechanisms underlying TTC estimation by humans.

In this study, we used mathematical modelling to explain how people estimate TTC for one object, and then extend the idea of understanding how people estimate TTC for two objects, simultaneously. Since our goal was to understand the mechanism of absolute TTC estimation, we used one and two-object PM tasks.

## 3. Materials and methods

Eighteen students from Amirkabir University of Technology-Tehran Polytechnic (9 women, 9 men, age 26.61 years±3.01 (mean±SD), min age 23, max age 33) took part in these tests voluntarily. All participants had normal or corrected-to-normal vision. They were healthy and without any known oculomotor abnormalities. All experiment protocols were approved by Iran University of Medical Sciences review board. Participants were naive with respect to the purpose of the experiment, gave written informed consent to their participation in the experiment, and were paid for their participation.

### 3.1. Apparatus and experimental procedure

Participants sat on a chair facing a 17″ computer display located at a viewing distance of approximately 50 cm, in a silent room with normal light. Stimuli were generated with MATLAB and presented on an Asus computer equipped with a 2.90 GHz Intel Corei7 processor. The screen resolution was 1920×1080 pixels (horizontal by vertical) and the display rate was 60 Hz.

### 3.2. Experiment 1 (one-ball experiment)

In the first section (hereafter referred to as “one-ball experiment”), time-to-contact (TTC) estimates for a green ball (diameter of 1 cm) moving at constant speed on a 25 cm × 12.5 cm (horizontal by vertical) frontoparallel plane from left to right were obtained using a prediction motion (PM) task (see [5]). The constant speeds were randomly selected from three values: 2 cm/s, 4 cm/s, 6 cm/s. After 1.5 seconds, the ball passed behind an 18 cm × 8 cm dark grey rectangle (hereafter referred to as “occluder”) that obscured its trajectory. A 0.3 cm × 8 cm vertically-oriented red line was shown at one of five different positions on top of the occluder (at a distance of 5.5 cm, 8 cm, 10.5 cm, 13 cm, 15.5 cm from the starting point of the occluder).

Participants pressed the spacebar key to start the test. After a delay of 2 s, the ball started to move at one of the above mentioned constant speeds, in a horizontal straight line towards the finish line. After 1.5 s, the ball passed behind the occluder and continued its motion to reach the finish line. The ball did not reappear after it was occluded. Participants had 10 seconds to press the “down” arrow key to indicate the instant at which they judged the ball would collide with the red finish line. No feedback on TTC estimation error was provided, but a smiley emoji was presented at the end of each trial if the individual finished the trial by pressing the down button and a sad emoji was presented if the subject did not press the down button in the 10-second interval and missed that trial. At the beginning of each trial we had a countdown from three to one, before the ball started moving. This caused a 2-second delay between two consecutive trials. Figure 1(a) shows the schematics of the one-ball experiment. In this experiment, each trial condition (combination of speed and end line position) was presented 10 times in random order, for a total of 150 trials.

**Figure 1.**
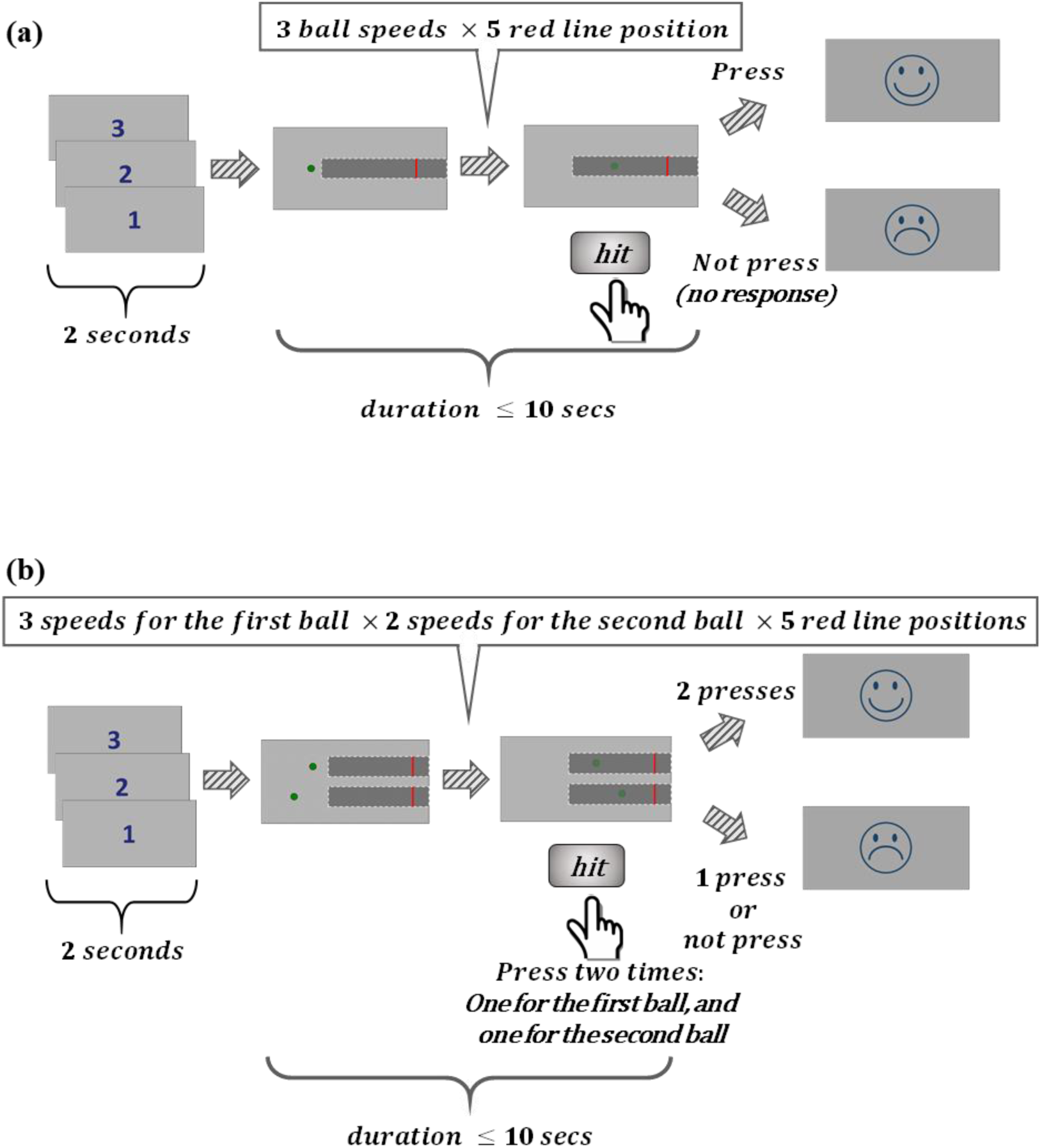
Experimental paradigm, (a) Schematic of one-ball experiment: after an ITI of 2 seconds, subject saw a rightward moving green ball that was presented 1.5 seconds before disappearing under an opaque cover. Subjects had to estimate the TTC for this ball with respect to the red line drawn over the cover by pressing a key after the ball went under the cover. If the subject pressed within 10 seconds a happy emoji would be presented for 0.5 seconds, otherwise, a sad emoji would be presented for 0.5 seconds. No other feedback about accuracy of TTC judgement would be provided, (b) Schematic of two-ball experiment: same procedure as the one-ball experiment except that two green balls with different speeds moving rightward were shown. Both balls went under the cover at the same time and subjects had to press the down button to estimate the TTC of each ball with respect to the red line.

### 3.3. Experiment 2 (two-ball experiment)

After completing the one-ball experiment, participants were tested in a second section (hereafter termed “two-ball experiment”), in which two balls were presented simultaneously and moved on parallel horizontal trajectories from left to right with different velocities. The initial point for each ball was determined so that (both balls) were visible for 1.5 seconds before going behind the occluder. Therefore, both balls started moving simultaneously and reached the cover simultaneously. TTC estimates were obtained using the same method as in the one-ball experiment. The number of conditions in the two-ball experiment was thirty (three velocities for one of the balls, two other velocities for the other ball, and five positions for the red finish line). Each condition was presented 5 times in random order in a session, resulting in the total number of 150 trials. Figure 1(b) shows the schematics of the two-ball experiment.

### 3.4. Mathematical background

We have used the following distributions for fitting TTC estimate for each subject:

1) Gaussian (normal) distribution [32]:

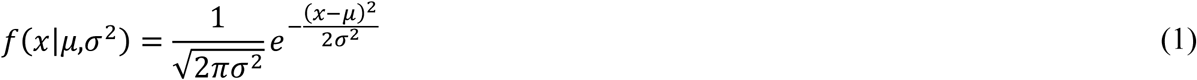

where *μ* is the mean or expectation of the distribution and *σ* is the standard deviation.
2) inverse Gaussian (IG) distribution [32]:

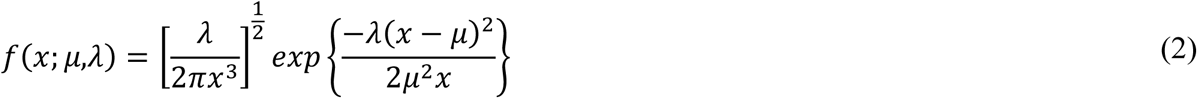

for *x* > 0. where *μ* > 0 is the mean and *λ* > 0 is the shape parameter.
3) Gamma distribution [32]:

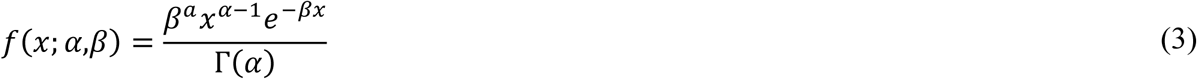

for *x* > 0 and *α,β* > 0, where Γ(*α*) is the complete Gamma function.
4) Weibull distribution [32]:

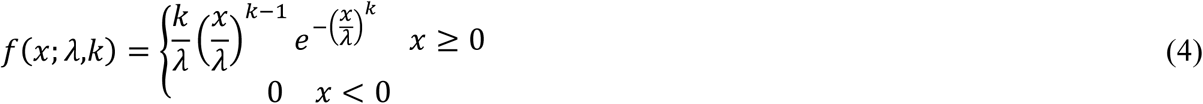

where *k* > 0 is the shape parameter and *λ* > 0 is the scale parameter of the distribution.
5) ex-Gaussian distribution [33]:

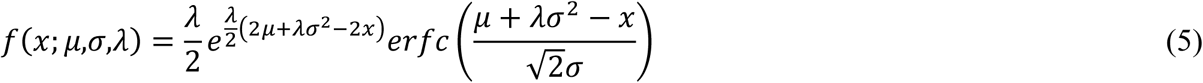

where *erfc* is the complementary error function defined as:

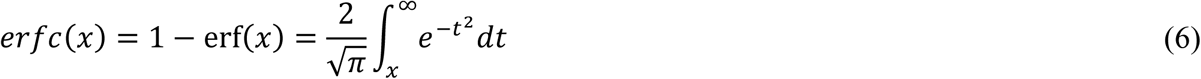

Note that all distributions considered except for ex-Gaussian have two free parameters thus R^2^ was fair with respect to the number of free parameters. The ex-Gaussian has three parameters so in theory it should have an advantage for fitting the data. To account for this difference in the number of parameters, we used the adjusted-R^2^ between the data cumulative distribution function (CDF) and predicted CDF using parameters fitted for each distribution as the arbitrator for the best fitting distribution. Ordinary *R*^*^2^*^ is defined as:

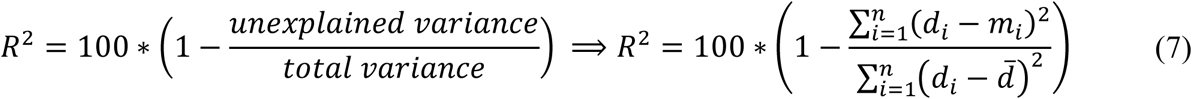

where *n* is the number of data points, *d*_*i*_ is the *i*th data point, *m*_*i*_ is the model fit for the *i*th data point, and *d* is the mean of the data points. Likewise, adjusted *R*^*^2^*^ which is a criteria to compare accuracy of models with different number of parameters, is defined as below:

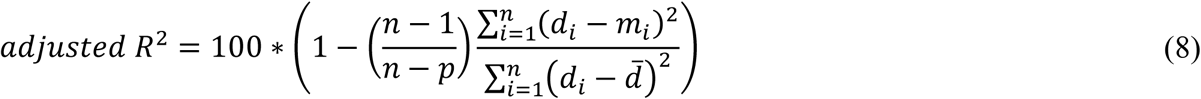

where *n* is the number of observations, and *p* is the number of parameters of the fitted model.

## Results

Figure l(a-b) shows the schematics of the one-ball and two-ball experiments. First, response times (RTs) of the one-ball experiment were studied to find out their general trend. The response times for trials belonging to the same condition were averaged (each trial was repeated 10 times randomly during the session) in the one-ball experiment and plotted in different colours for each participant in Figure 2(a).

**Figure 2.**
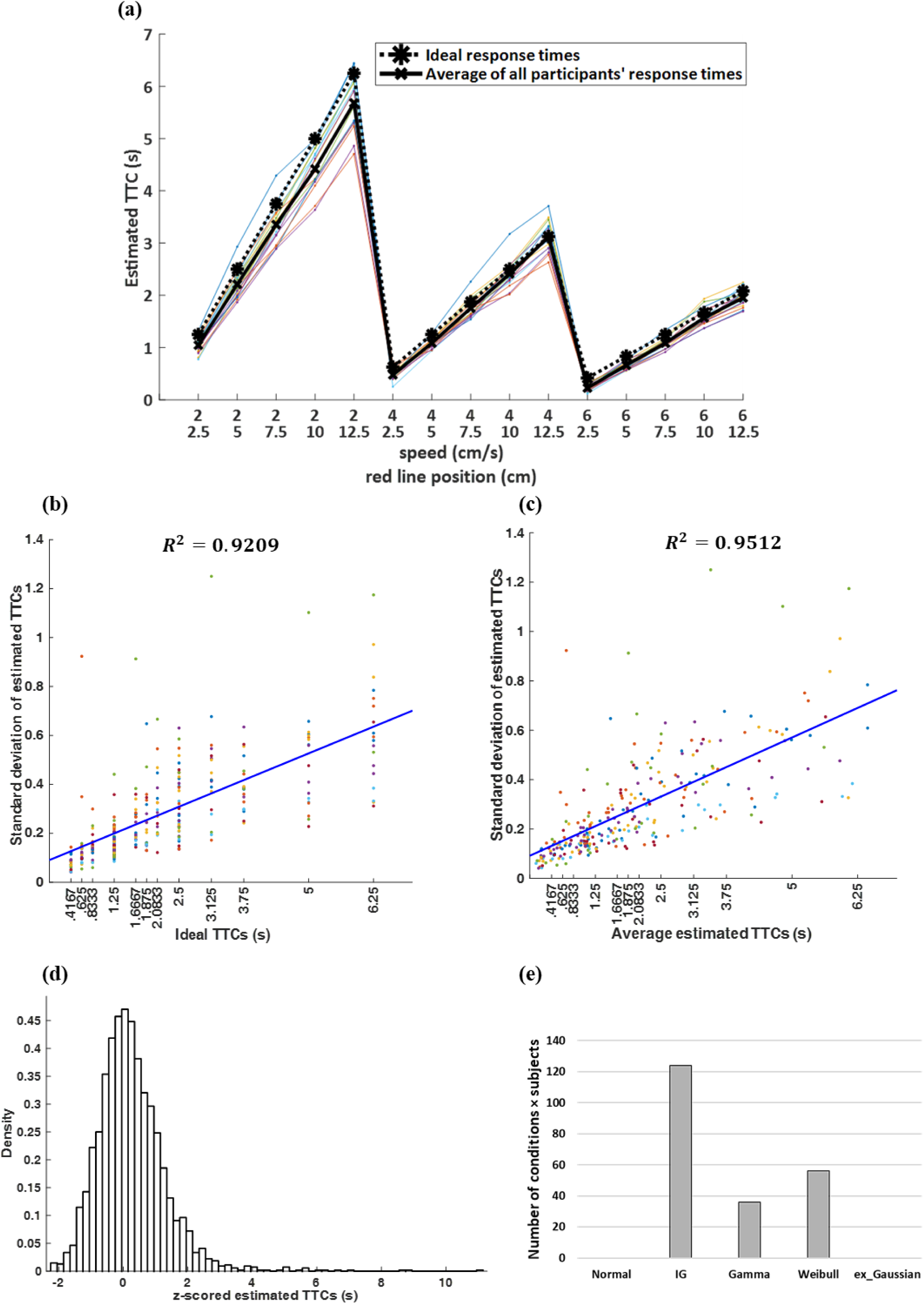
TTC estimates and error patterns in one-ball experiment, (a) Estimated TTCs for one-ball experiment. Each marker on the horizontal axis represents a pair composed of ball speed and red line position: (ball speed (cm/s), red line position (cm)), and the vertical axis shows the response times averaged on similar trials. The dashed black plot shows the ideal TTC, and the thick black line shows the average of all subjects’ responses (subject’s estimated TTC). The coloured plots show the averaged response times related to similar trials (each trial was repeated 10 times randomly) for each participant, (b) Estimated TTC variability versus ideal TTCs for the one-ball experiment, (c) Response times variability versus average estimated TTC for each subject for the one-ball experiment. Each dot belongs to one participant in a given condition. Each colour is for one participant, (d) Aggregate density of z-scored estimated TTCs for all subjects in all trials and five probability distributions fitted to it. (e) Number of conditions × subjects in which each of the five probability distributions provided the best fit by maximizing the likelihood under that distribution.

Results in Figure 2(a) show that the subjects’ estimated TTCs are close to the ideal TTCs, but on average, subjects tend to slightly underestimate TTCs (thick black line below dashed black line). In addition, for each speed, increasing the distance of the finish line from the start point, resulted in an increase in the variability of the responses among subjects (more spread in coloured lines). In order to better study this variability, the standard deviations of response times are plotted in Figure 2(b-c). Figure 2(b), shows that the TTC estimate variability increases monotieally for longer ideal TTCs. Indeed, this monotonic increase can be well explained in linear fashion (Figure 2(b) blue line, R2=0.92). As mentioned previously, participants tended to underestimate the TTCs (Figure 2(a)). Therefore, we hypothesized that if the variability in TTC is to increase linearly with time, this relationship should be stronger if one considers the variability as a function of subjects estimated TTCs rather than the ideal TTCs. Indeed, the linear fit between TTC variability and estimated TTC resulted in a modest increase in the goodness of fit (Figure 2(c) blue line, R2=0.95). To further characterize response variability, we looked at the probability density function of response times. Figure 2(d) shows the density of response times in all conditions and for all participants. For this figure, in each condition we collected all the data for all the participants. Then we normalized estimated TTCs by z-scoring the data (mean value of zero and variance of one). It can be seen that the distribution of estimated TTCs is rightward-skewed.

To determine the distribution underlying the response variability, the estimated TTCs in each condition for every participant (10 similar trials are considered as one condition) were fit separately to each multiple candidate distribution functions (see methods). Then the adjusted *R*^*^2^*^ (see methods) was computed for each condition (each condition is a combination of the speed of the ball and the position of the red line) and each participant. Next, the number of conditions for which each of the five probability distributions were the best fit (has maximum adjusted *R*^*^2^*^) were counted across all conditions and all participants and were compared across the distributions (Figure 2(e)). As can be seen inverse Gaussian distribution provided a better fit to the data in an overwhelming majority of conditions and subjects.

This fact that the density of estimated TTCs is best fit by an inverse Gaussian, together with the fact that standard deviation of the responses showed an increase with increasing the TTC, led us to propose a drift-diffusion model to explain the cognitive process underlying the TTC estimation in our human subjects. Figure 3(a) shows the schematic of this model for one object. In this model we assume that subjects track passage of time after each small time step Δt (similar to pacemaker accumulator model) [17, 34-37]) with an additive Gaussian noise at each time step. Such a system can be modelled with a stochastic process with the following formulation:

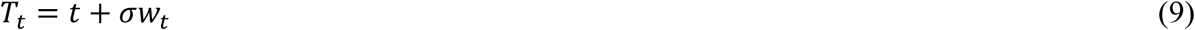

where *T*_*t*_ indicates the subject’s estimated time (decision variable) and the decision thresholds being *P/v*, where *P* is the position of the finish line, and *v* is the velocity of the ball. The right side of equation (9) which denotes the noisy evidence obtained during the trial, is composed of two terms: a constant drift *t*, that is the actual time, and the diffusion term *σw*_*t*_, which represents white noise drawn from a Gaussian distribution, *σ* is the standard deviation of the integration noise, and *w*_*t*_ is a Brownian motion. If the decision bounds are ignored, *T*_*t*_ is normally distributed with probability density 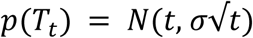 [38]. Consequently, the variance across trials of the temporal evolution of *T*_*t*_ increases with *t.* However, it can be shown that the time to hit a decision threshold in such a drift-diffusion model is rightward-skewed and follows an inverse Gaussian distribution.

**Figure 3.**
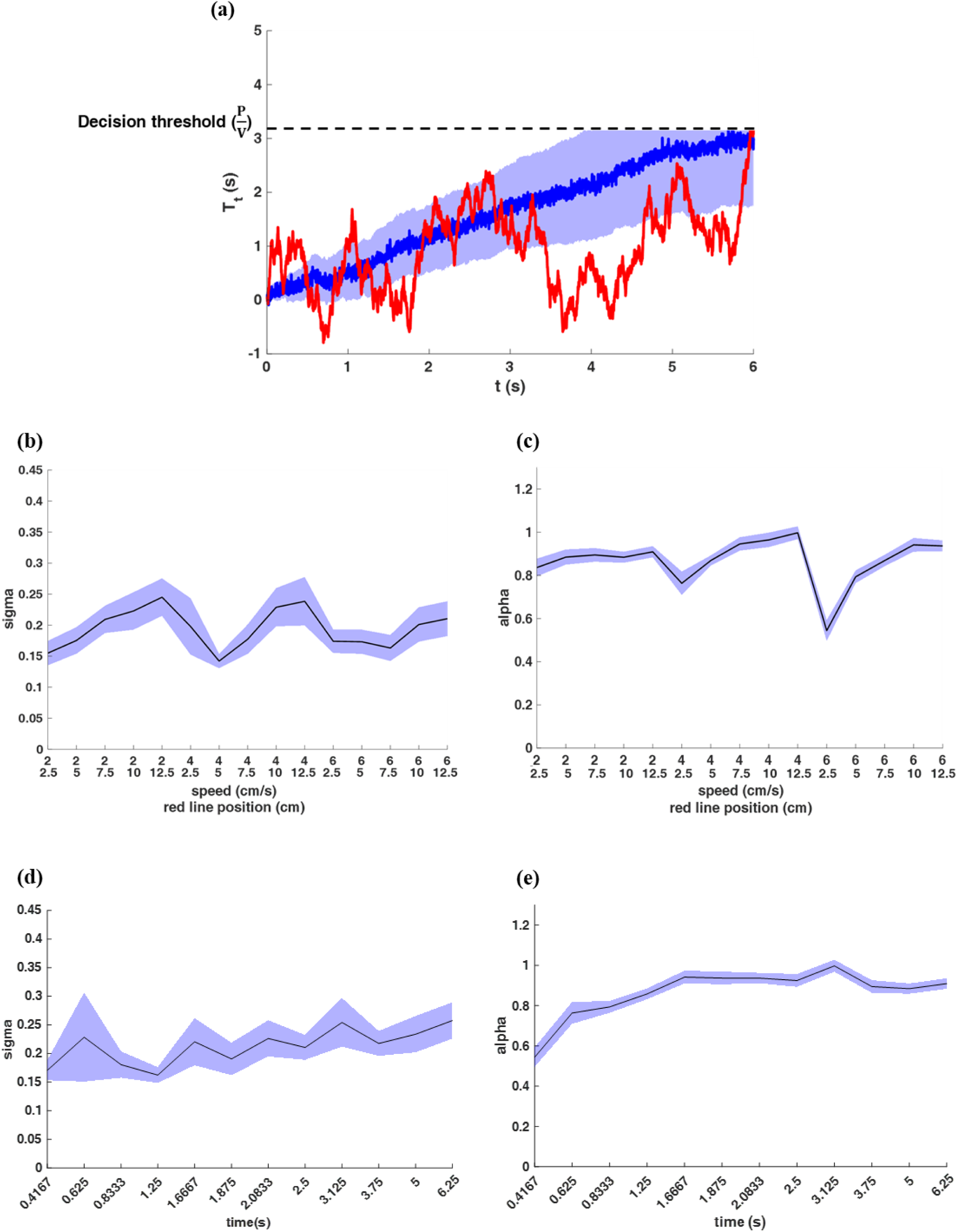
A biased drift-diffusion model can explain the TTC estimates, (a) Schematic of the drift-diffusion model for one object. Red plot indicates decision variable, blue plot indicates the mean of 100 drift-diffusion models, and the red shade around it is the variance of this plot. The horizontal dashed line shows the decision threshold. When a decision variable reaches this threshold, a decision will be made, (b) *σ* values for all trials in the one-ball experiment sorted based on different trial conditions, (c) *σ* values for all trials in the one-ball experiment sorted based on ideal TTCs. (d) *α* values for all trials in the one-ball experiment sorted based on ideal TTCs. (e) *α* values for all trials in the one-ball experiment sorted based on ideal TTCs.

Therefore, drift-diffusion model could be a good candidate for TTC estimation. For one-ball experiment we have:

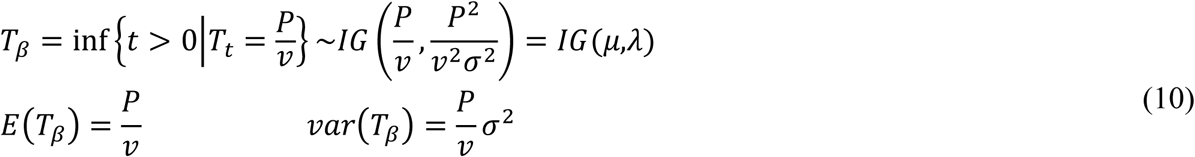

where *T*_*β*_ indicates the time to hit the threshold (first passage time), and *IG* indicates the inverse Gaussian model. *P* is the position of the finish line, and *v* is the speed of the ball.

Thus the integration noise *σ* can be obtained from the expected value and variance of the estimated TTC:

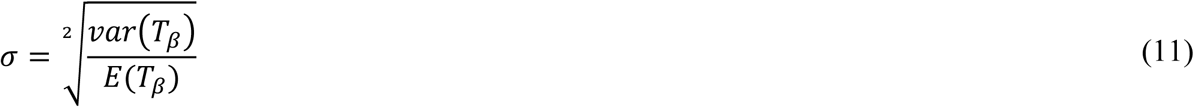

As mentioned above (see Figure 2 (a)), participants tend to underestimate the TTCs. To account for this observation, we assume that subjects use a scaled decision threshold such that:

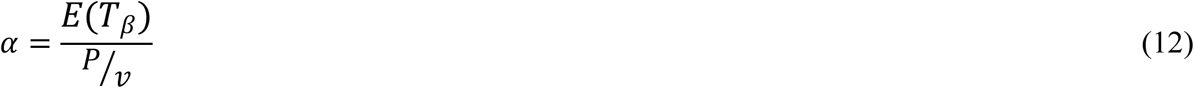

Thus, the full stochastic process will have the following formulation:

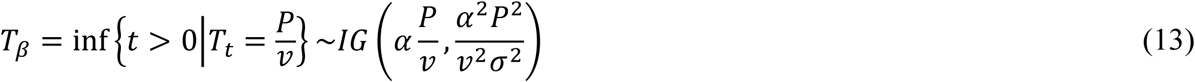

Figure 3(b-e) show the fitted values of *σ* and *α* for the one-ball experiment for each condition (each condition is a combination of the speed of the ball and the position of the red line) across all subjects. Results show that the effect of condition on both *σ* and *α* was significant (F(14,269)=2.032, *p*=0.016 for *σ,* and *F*(14,269)=20.532, *p*<0.001 for *α).* The integration noise seemed to increase at higher speeds while TTC underestimation seemed lower at higher speeds (i.e. *α* closer to 1 in higher speeds, Figure 3 (c & e)). Furthermore, values of *σ* and *α* were significantly different across subjects (F(17,268)=8.153, *p*<0.001 for *σ*, and *F*(17,268)= 1.691, *p=*0.045 for *α*) suggesting that subjects had different levels of integration noise and TTC underestimation.

Next to see whether subjects congnitive model using drift-diffusion in one-ball experiment can extend to cases with more balls, we repeated the analysis in the two-ball experiment. Figure 4(a & b) shows the response times of the first ball and the second ball in the two-ball experiment. Once again subjects’ estimated TTC tended to be close to the ideal TTCs. However, there was a small tendency to overestimate TTC of the first ball (ball with shorter TTCs) and the second ball in the speed of 4 cm/s (shorter TTCs compared to the speed of 2 cm/s). Also, there was a tendency to underestimate the TTC of the second ball in the speed of 2 cm/s. Indeed the average TTC error across conditions and subjects was significantly different between the first and second ball in the two-balls experiment (*F*(9,2520)=6.026, *p*<0.001 for different conditions, and *F*(17,2520)=26.652, *p*<0.001 for different subjects for the first ball; and *F*(9,2520)=73.558, *p*<0.001 for different conditions, and *F*(17,2520)=20.472, *p*<0.001 for different subjects for the first ball). The error in the second ball was slighlty negative on average and was similar to the negative error seen in the one-ball experiment (time underestimation).

**Figure 4.**
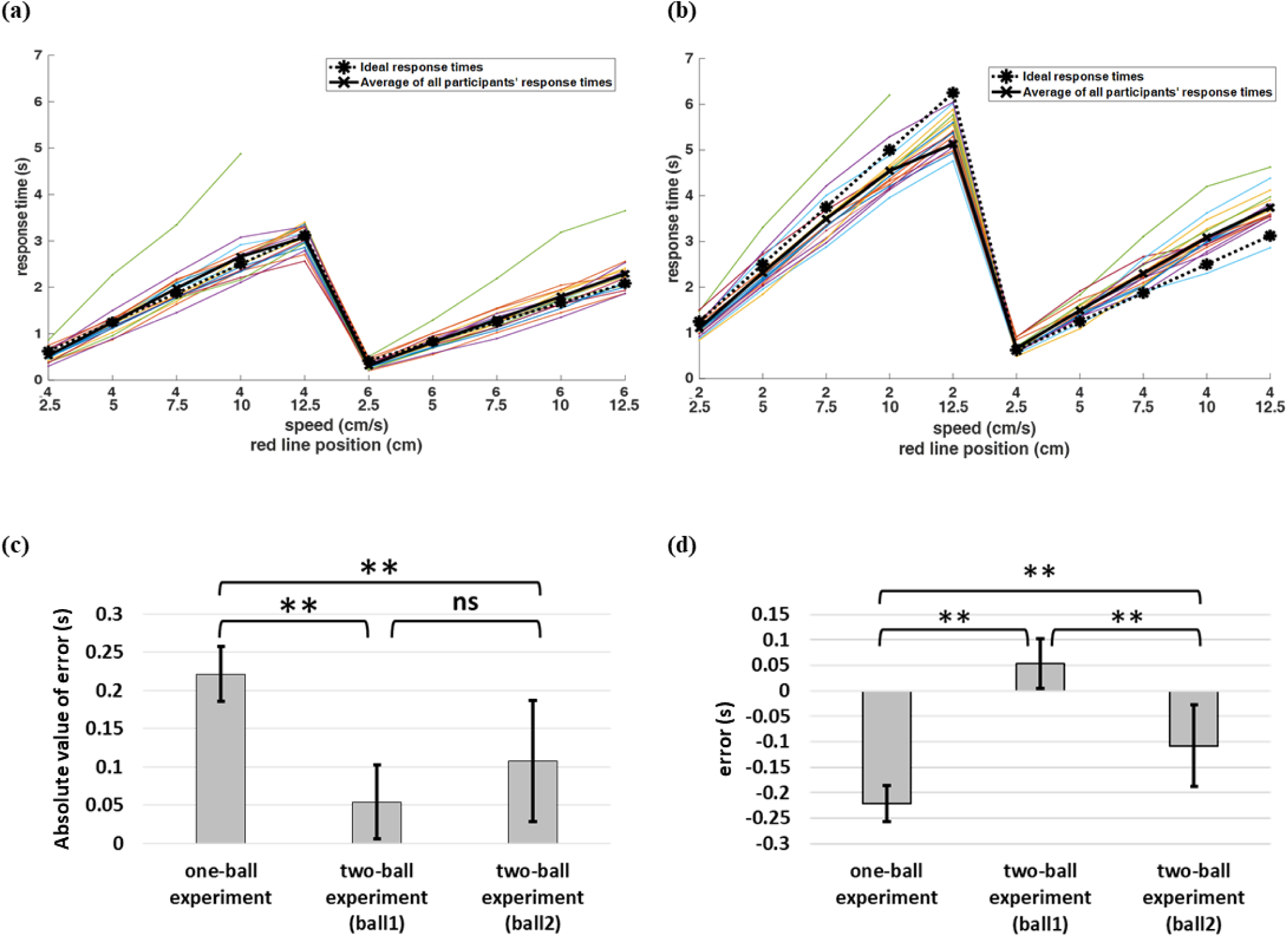
TTC estimates and error patterns in the two-ball experiment (a) Estimated TTCs for the first ball in the two-ball experiment, (b) Response times for the second ball in the two-ball experiment. Each marker on the horizontal axis represents a pair composed of ball speed and red line position: (ball speed (cm/s), red line position (cm)), and the vertical axis shows the estimated TTCs averaged on similar trials. The dashed black plot shows the ideal TTC judgments, and the thick black line shows the average of all subjects’ responses, (c) Absolute average TTC error for one-ball and two-ball experiments. Error bars throughout this paper show the standard error of mean (SEM). (d) Average TTC error for one-ball and two-ball experiments.

Figure 4 (c-d) shows the mean of error values of all trials and all participants for the one-ball experiment and the first ball and the second ball in the two-ball experiment. While the error in TTC estimation was in general low, the absolute error seemed to be slightly larger in the one-ball experiment (*F*(2,8099)=275.426, *p*<0.001, post hocs between one-ball experiment and the first ball *p*<0.001 and the second ball p=0.006). The absolute error was not significantly different between the first and second ball (p=0.015). However the direction of error was toward a slight overestimation in the first ball and a slight underestimation in the second ball (*F*(2,8099)= 166.283, *p*<0.001, all post hocs *p*<0.001).

In the two-ball experiment, the distribution of estimated TTC again showed features consistent with an underlying drift-diffusion process, that is: 1) the variance of estimated TTC increased linearly with ideal (Figure 5(a & b) *R*^2^=0.3861 for first ball, *R*^2^=0.414 for second ball) and average TTCs (Figure 5(c & d) *R*^2^=0.5244 for first ball, *R*^2^ =0.5131 for second ball) and 2) the shape of estimated TTC distribution was best described in majority of conditions across all subjects with inverse Gaussian for both the first and second ball (Figure 5(e & f)).

**Figure 5.**
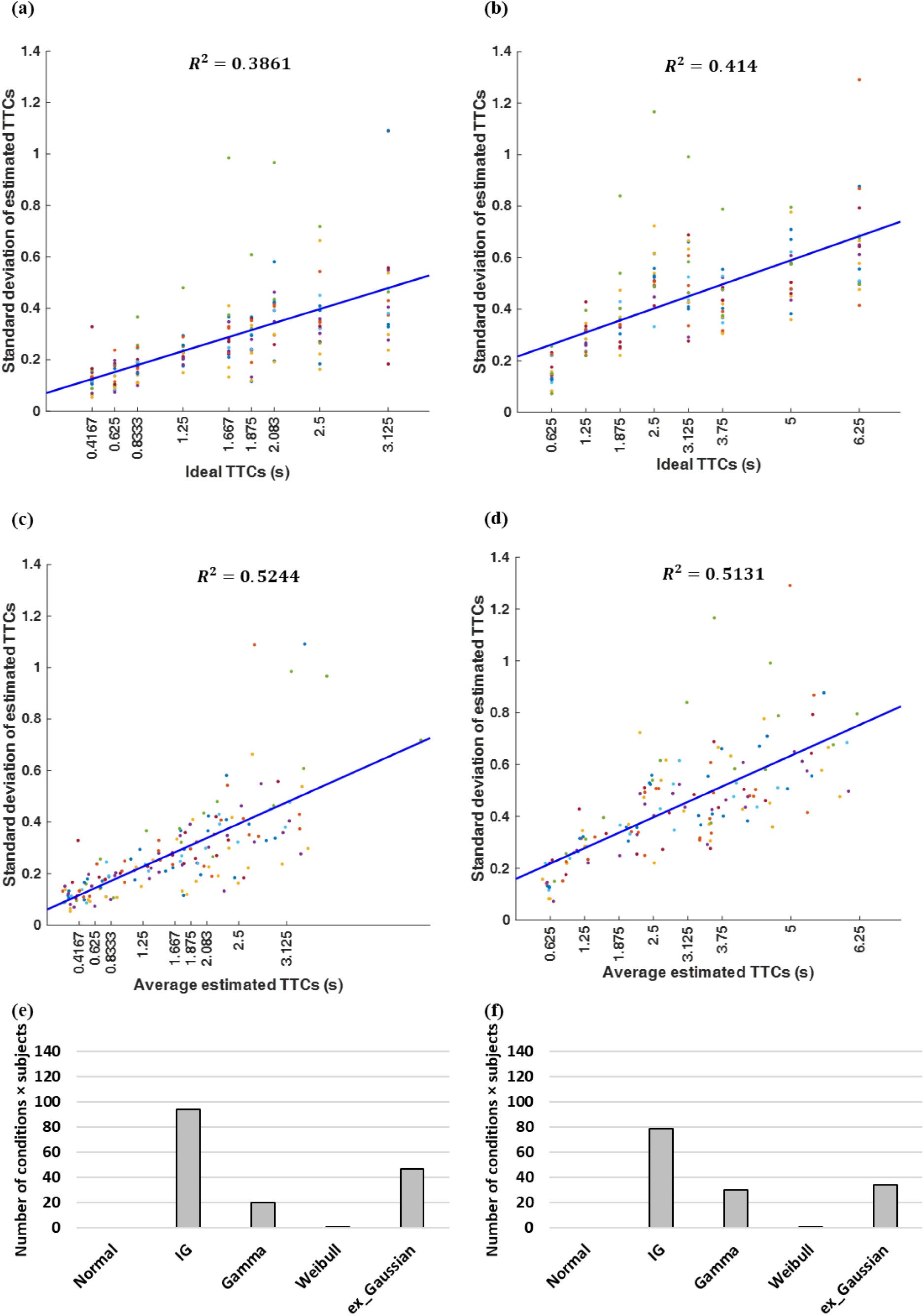
(a) Estimated TTC variability versus ideal TTCs for the first ball of the two-ball experiment, (b) Response times variability versus average estimated TTC for each subject for the first ball of the two-ball experiment, (c) Estimated TTC variability versus ideal TTCs for the second ball of the two-ball experiment, (d) Response times variability versus average estimated TTC for each subject for the second ball of the two-ball experiment. Each dot belongs to one participant in a given condition. Each colour is for one participant.

Theoretically, tracking objects in the two-ball experiment could be done either with a single decision variable or with two decision variables that evolve independent from each other (Figure 6(a & b)). The formulation of drift-diffusion process in the case of single DV is:

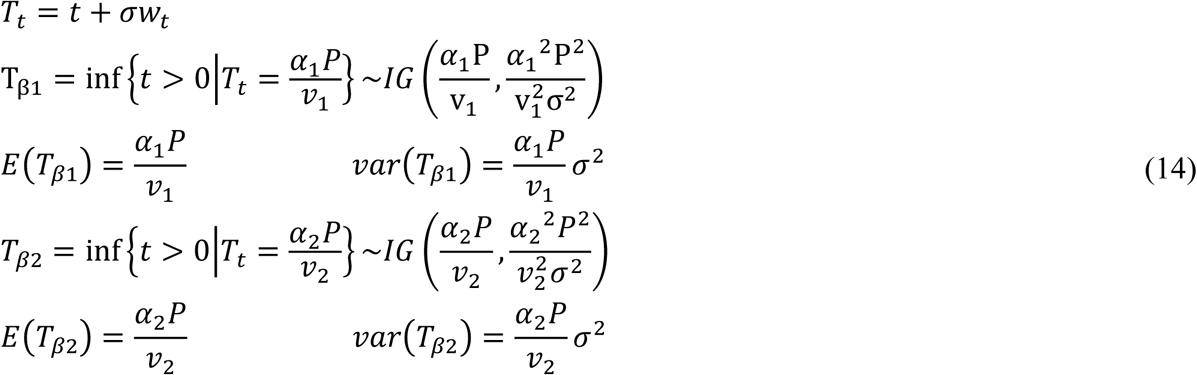

where *T*_*t*_ describes the drift-diffusion model for both balls, but the threshold to make a decision for each ball is different. The threshold for the first ball is 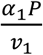 and for the second ball 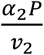, where *v*_1_ and *v*_2_ are respectively the speed of the first ball and the second ball. And *P* shows the position of the red finish line for both balls. *IG* indicates the inverse Gaussian model.

**Figure 6.**
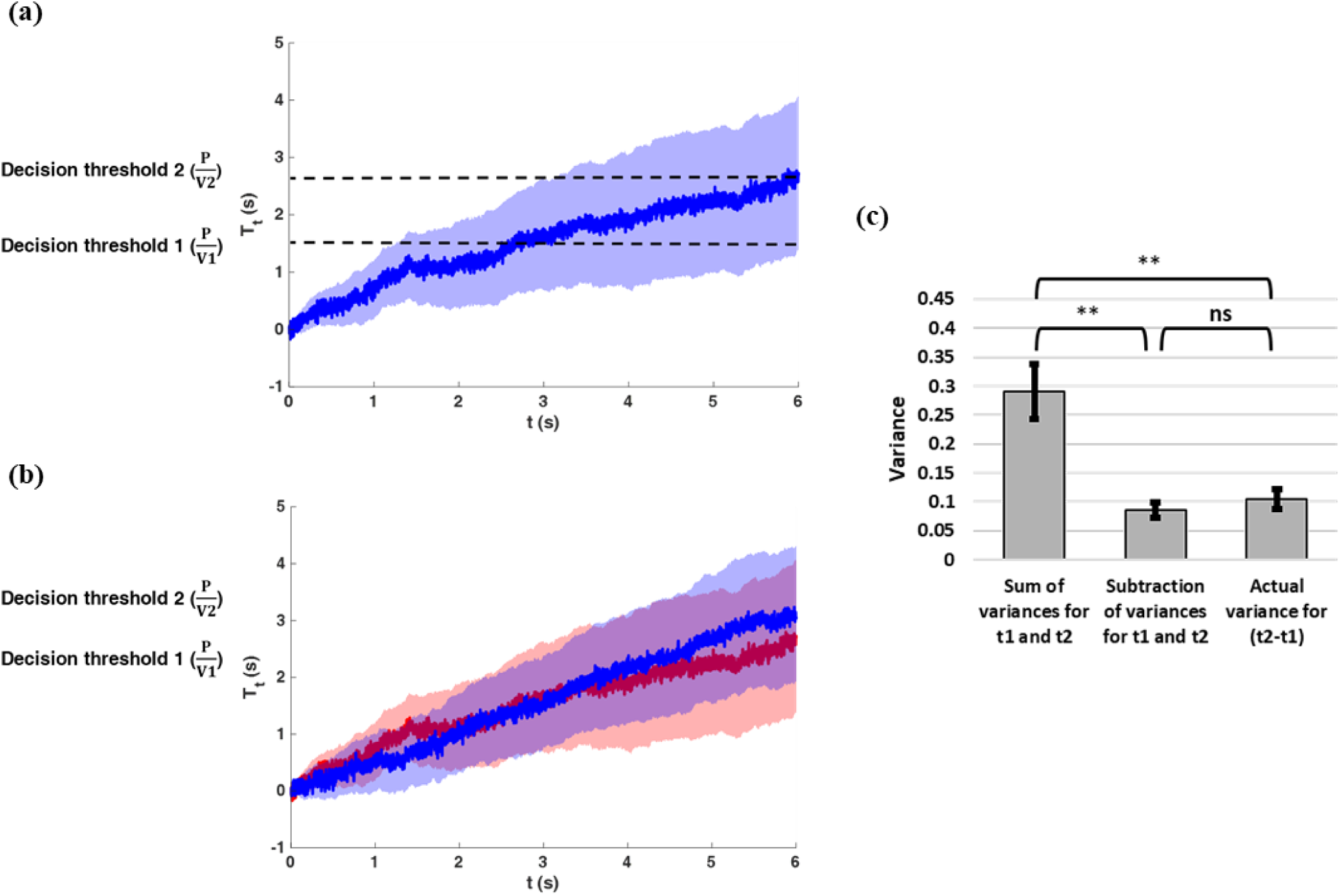
Two alternative drift-diffusion models can be used for TTC of two balls, (a) Schematic of the drift-diffusion model for two objects, when the participant considers a single decision variable for both balls. Here, the participant estimates the TTC of the first ball when the decision variable hits the first decision threshold and TTC of the second ball when the same decision variable now hits the second decision threshold, (b) Schematic of the drift-diffusion model for two objects, when the participant considers a separate decision variable for each ball. Here, the participant estimates TTC of the first ball when the first decision variable hits the first decision threshold and TTC of the second ball when the second decision variable hits the second decision threshold. (c)The variance of difference in TTC estimate for ball one (t2) and ball two (t2) in two ball experiment is consistent with a drift-diffusion model with a single decision variable.

While for two independent DVs one can write:

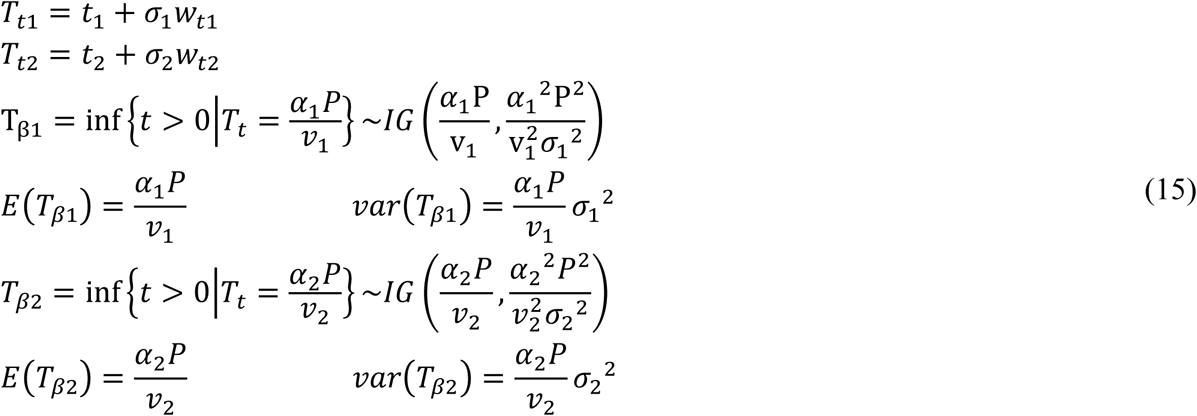

where *T*_*t*1_ and *T*_*t*2_ indicate respectively the drift-diffusion model for the first ball and the second ball.

To know whether a single or double process was used by our subjects, one can look at the relationship between the errors of estimated TTC between the two balls. The prediction is that if for subjects used a single DV, the error in the estimated TTC of the first and second ball would be correlated. That is if one were to underestimate the first ball he/she would be more likely to underestimate the second ball and vice versa. But in the two DV condition the errors would be uncorrelated.

Thus, if one were to form the variance of subtracted estimated TTCs for the first and second ball in the single DV condition, one would have:

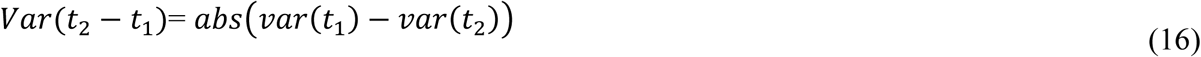

and in the double DV condition, one would have:

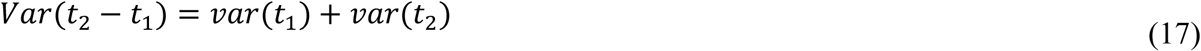

Figure 6(c) shows the values of actual variance of the t1-t2 subtraction, and the subtraction and sum of t1 and t2 variances for all participants, and all trials. Interestingly, results show that the variance of subtraction is significantly different from the sum of variances (*F*(2,53)=14.039, *p*<0.001, post hoc *p*<0.001) and instead is very close to subtraction of variances (post hoc *p=*0.66). This means that on average participants tended to use a single DV for both balls in the two-ball experiment.

*σ* and *α* values for the two-ball experiment are calculated and shown in Figure 7. Figure 7(a & b) show the *σ* values for the first ball and the second ball, and Figure 7(c & d) show the *α* values for the first ball and the second ball in the two-ball experiment. Both *σ* and *α* values seemed to be higher in the two-ball compared to one-ball experiment. Also, for the second ball in the two-ball experiment there is a small increase in these values compared to the first ball in the two-ball experiment. Results show that the effect of condition on *σ* and *α* for both balls was significant (*F*(8,161)= 1.746, *p=*0.092 for *σ* of the first ball, and *F*(8,161)=5.080, p<0.001 for *α* of the first ball and *F*(7,143)= 2.142, *p*=0.043 for *σ* of the first ball, and *F*(7,143)=5.798, *p*<0.001 for *α* of the second ball).

**Figure 7.**
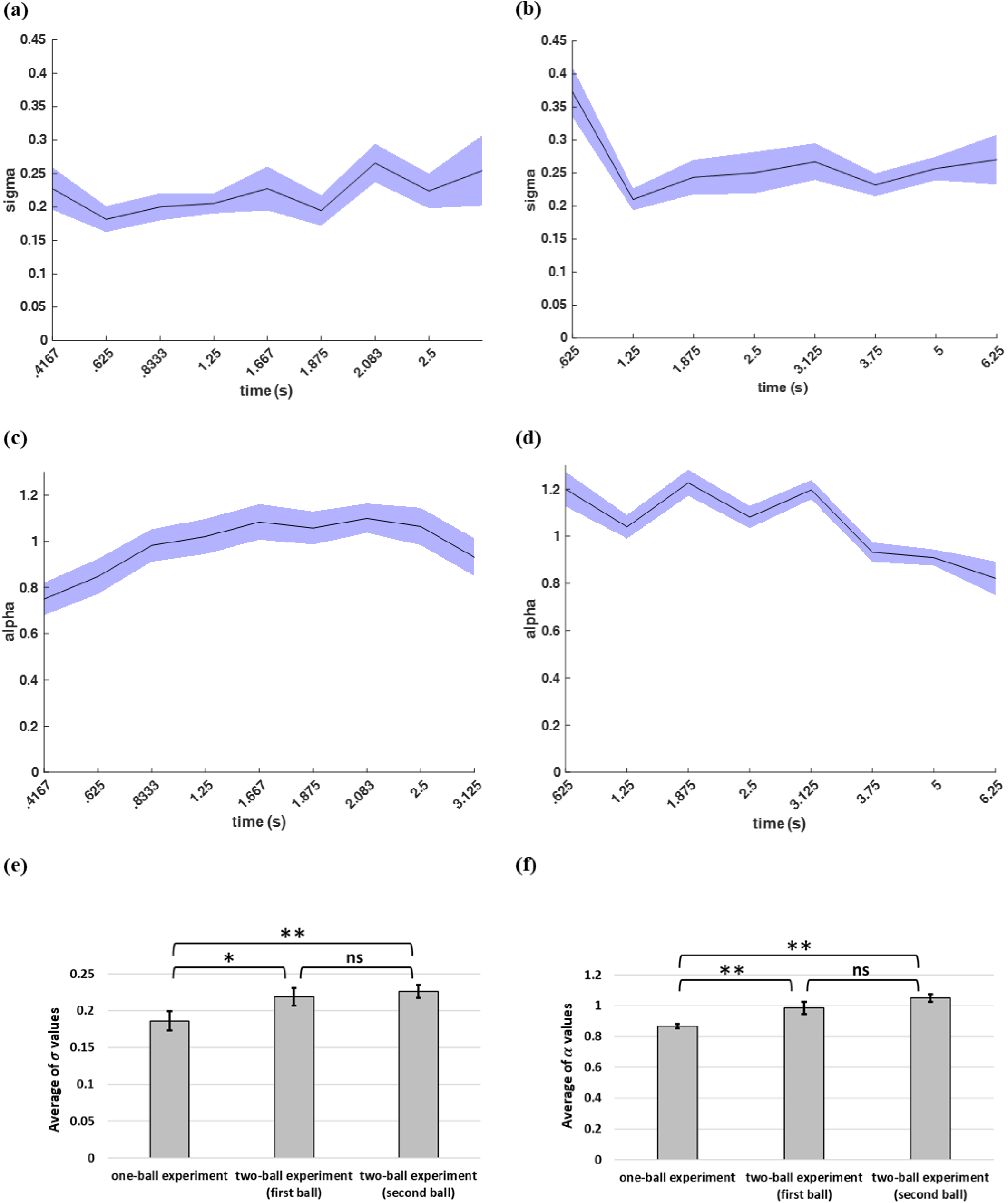
(a) *σ* values for the first ball in all trials of the two-ball experiment, (b) *σ* values for the second ball in all trials of the two-ball experiment, (c) *α* values for the first ball in all trials of the two-ball experiment, (d) *α* values for the second ball in all trials of the two-ball experiment, (e) average of *σ* values, and (f) average of *α* values for one-ball and two-ball experiments.

Figure 7 (e & f) show the average of *σ* and *α* values for all participants and all trials in one-ball experiment and two-ball experiment. ANOVA for *σ* values showed that there was a significant difference between *σ* values (*F*(2,53)=5.132, *p=*0.009, post hoc one-ball with first ball *p*=0.026 and second ball *p*=0.003). But, the difference between the first ball and the second ball in the two-ball experiment was not significant *(p=*0. 439). Also, ANOVA for *α* values showed that there was a significant difference between *α* values (*F*(2,53)=10.615, *p*<0.001, post hoc one-ball with first ball *p*=0.005 and second ball *p*<0.001). Again the difference between the first ball and the second ball in the two-ball experiment was not significant (*p*=0.105). The fact that that *σ* and *σ* values for the first and second ball were similar is consistent with and is a further confirmation of the fact that a single DV was used to track both balls.

## 4. Discussion

We wanted to construct a cognitive model for human subjects tracking objects in a prediction motion task. We asked human subjects to estimate TTC of a ball hitting an end line after it disappeared from sight. Our results based on pattern of TTC estimates showed that variance of TTC error scaled with absolute TTC estimate and followed a rightward-skewed distribution that was best fit by an inverse Gaussian distribution (Figure 2(d-e)). These findings suggest that the subjects’ temporal estimates could be generated by a drift-diffusion decision variable (Figure 3(a)).

Overall subjects tended to underestimate the actual TTC consistent with a biased drift-diffusion model. The integration noise for the decision variable was relatively stable across speeds and end line positions (Figure 3(b)). When tracking two balls simultaneously subjects’ average performance was not much worse compared to one-ball condition with the bias and integration noise being comparable in the one-ball and two-ball condition (Figure 3(d-e) and Figure 7(a-d)). Interestingly, analysis of pattern of TTC errors for the two-ball experiment showed that subjects were using a single decision variable for tracking both balls, resulting in dependence between TTC estimates for the first and the second ball. This finding suggests that the errors in estimating the TTC for the first ball can easily propagate to cause miscalculation of the second ball TTCs. Our finding thus explains previous results on how errors on leading objects can affect errors for trailing objects [11, 28, 29, 31]. It should be mentioned that as in our two-ball experiment both balls went behind the occluder at the same time, it was possible for subjects to form a single DV for two balls. Whether the same strategy can be used for cases where two balls are to disappear at different times, needs to be tested in future experiments.

In the one-ball experiment, increasing the ideal TTC caused *α* values to increase for the short TTCs, saturating at about *α* = 1 for longer durations (Figure 3 (e)). This indicates that on average subjects tended to respond faster than they should in short but not long TTCs in the one-ball experiment. While the reason for this dependence is not clear at the moment this may be related to subjects effort not miss short TTCs. Such underestimation of time is also seen for the faster ball with the closest end line positions as well (Figure 7(c)). Although trends of changes in alpha between the two balls were not always easy to interpret (Figure 7(d)). Also, increasing the ideal TTCs caused a slight increment in the *σ* values in both one-ball and two-ball experiments (Figure 3 (d) and Figure 7(a-b)). These changes in *σ* values, although small, may indicate higher order effects that are not sufficiently modeled by a simple biased drift-diffusion model. The underlying mechanism for such changes in not currently known and needs to be investigated in future experiments.

Our finding indicates that subjects can be surprisingly accurate on average in their TTC estimates in both one and two object tracking conditions (Figure 2(a) and Figure 4(a-b)) despite the fact that we never provided feedback about the accuracy of TTC estimates to our subjects. We found a slight overall underestimation of TTC times in one-ball experiment (but not necessarily in two-ball experiment). This result is somewhat different from some of the previous literature which found larger average errors in TTC ([28, 29, 31]) (Figure 4(c)). Other studies reported small errors consistent with our results [39-42]). One possible reason for low bias in our task was that the time interval for the balls to move behind the occluder was 1.5 seconds which is about twice that of the some other studies (0.8 seconds in [28] and 0.8 seconds in [29]). Therefore, the participants in our experiment had enough time to accurately estimate the speed the balls before they disappeared. In addition, it is previously shown that TTC estimation outside the time interval of (0.4 seconds to 10 seconds) is not reliable[40]. That is because at least about 0.4 seconds is needed to push a button and report the estimated response and on the other hand, participants are not able to hold the model of object’s movement in their memory more than 10 seconds. The ideal TTCs in our experiment are within that time interval. Our subjects might have been further helped by the fact that the time under occluder was comparable to the visible time before the occlude a factor that is shown to facilitate temporal estimation [7, 43].

Some studies have shown that TTC estimates for each condition tends to be biased toward the mean TTC across all conditions [4, 6, 7, 44]. We did not observe such pattern in our experiments. One likely reason for this difference could be due to lack of feedback in our experiment. Such feedback can result in forming priors for the subjects centred around the mean TTC which would bias estimates toward the mean in each trial. Furthermore, the overall underestimation of TTCs similar to ours have been reported previously [8, 40].

In summary, we provided a rigorous cognitive model for object tracking based on a biased drift-diffusion model. Our model can predict the pattern of TTC errors and can provide a mechanism for explaining the dependence of errors when tracking two objects simultaneously. The fact that two objects were tracked with a single decision variable can show a serious limitation in working memory capacity for tracking two objects. Such a limitation can have catastrophic consequences for human observers in dealing with their environment for instance when trying to estimate TTC of incoming vehicles as miscalculation for object can adversely affect TTC estimate for other objects. Future experiments should address whether this limitation generalizes to other configurations (e.g. balls moving in orthogonal directions) and with more objects to track. Furthermore, the neural basis for tracking objects and its overlap or dissociation from areas involved in pure time-lapse estimate needs to be studied.

## Acknowledgements

We would like to thank so and so for their help with this study and helpful discussions. This work was supported by grant #.

## Author contributions

AD, HA, AG and FT designed the experiment. AD collected the data under HA supervision. AG proposed the drift diffusion model and designed the analysis and AD performed the analysis. AD and AG wrote the paper with input from HA and FT.

## References

1. Hecht, H. and G.J. Savelsbergh, Time-to-contact. Vol. 135. 2004: Elsevier.

2. Schiff, W. and R. Oldak, Accuracy of judging time to arrival: effects of modality, trajectory, and gender. Journal of Experimental Psychology: Human Perception and Performance, 1990. 16(2): p. 303.

3. Kaiser, M.K. and A.V. Phatak, Things that go bump in the light: On the optical specification of contact severity. Journal of Experimental Psychology: Human Perception and Performance, 1993. 19(1): p. 194.

4. Schiff, W. and M.L. Detwiler, Information used in judging impending collision. Perception, 1979. 8(6): p. 647–658.

5. Tresilian, J., Perceptual and cognitive processes in time-to-contact estimation: Analysis of prediction-motion and relative judgment tasks. Perception & Psychophysics, 1995. 57(2): p. 231–245.

6. Ahrens, M.B. and M. Sahani, Observers exploit stochastic models of sensory change to help judge the passage of time. Current Biology, 2011. 21(3): p. 200–206.

7. Chang, C.-J. and M. Jazayeri, Integration of speed and time for estimating time to contact. Proceedings of the National Academy of Sciences, 2018. 115(12): p. E2879–E2887.

8. Steeves, J., et al., Accuracy of estimating time to collision using only monocular information in unilaterally enucleated observers and monocularly viewing normal controls. Vision research, 2000. 40(27): p. 3783– 3789.

9. de la Malla, C. and J. López-Moliner, Hitting moving targets with a continuously changing temporal window. Experimental brain research, 2015. 233(9): p. 2507–2515.

10. Cavallo, V. and M. Laurent, Visual information and skill level in time-to-collision estimation. Perception, 1988. 17(5): p. 623–632.

11. Hecht, H. and G.J. Savelsbergh, Theories of time-to-contact judgment, in Advances in psychology. 2004, Elsevier. p. 1–11.

12. López-Moliner, J., D.T. Field, and J.P. Wann, Interceptive timing: Prior knowledge matters. Journal of Vision, 2007. 7(13): p. 11–11.

13. Smeets, J.B., et al., Is judging time-to-contact based on ‘tau’? Perception, 1996. 25(5): p. 583–590.

14. Lee, D., et al., Visual timing in hitting an accelerating ball. The Quarterly Journal of Experimental Psychology, 1983. 35(2): p. 333–346.

15. Allan, L.G., The perception of time. Perception & Psychophysics, 1979. 26(5): p. 340–354.

16. Brown, S.W., Time perception and attention: The effects of prospective versus retrospective paradigms and task demands on perceived duration. Perception & Psychophysics, 1985. 38(2): p. 115–124.

17. Buhusi, C.V. and W.H. Meck, What makes us tick? Functional and neural mechanisms of interval timing. Nature Reviews Neuroscience, 2005. 6(10): p. 755.

18. Diedrichsen, J., S.E. Criscimagna-Hemminger, and R. Shadmehr, Dissociating timing and coordination as functions of the cerebellum. Journal of Neuroscience, 2007. 27(23): p. 6291–6301.

19. Eagleman, D.M., Human time perception and its illusions. Current opinion in neurobiology, 2008. 18(2): p. 131–136.

20. Grondin, S., Timing and time perception: a review of recent behavioral and neuroscience findings and theoretical directions. Attention, Perception, & Psychophysics, 2010. 72(3): p. 561–582.

21. Kanai, R., et al., Time dilation in dynamic visual display. Journal of vision, 2006. 6(12): p. 8–8.

22. Meck, W.H., Neuropsychology of timing and time perception. Brain and cognition, 2005. 58(1): p. 1–8.

23. Merchant, H., et al., Interval tuning in the primate medial premotor cortex as a general timing mechanism. Journal of Neuroscience, 2013. 33(21): p. 9082–9096.

24. Pöppel, E., Time perception, in Perception. 1978, Springer. p. 713–729.

25. Novak, J.L.B., Judgments of absolute time-to-contact in multiple object displays: Evaluating the role of cognitive processes in arrival-time judgements. 1997, Texas Tech University.

26. McLeod, R.W. and H.E. Ross, Optic-flow and cognitive factors in time-to-collision estimates. Perception, 1983. 12(4): p. 417–423.

27. Bootsma, R.J. and R.R. Oudejans, Visual information about time-to-collision between two objects. Journal of experimental psychology: human perception and performance, 1993. 19(5): p. 1041.

28. Baurès, R., D. Oberfeld, and H. Hecht, Judging the contact-times of multiple objects: Evidence for asymmetric interference. Acta Psychologica, 2010. 134(3): p. 363–371.

29. Baurès, R., D. Oberfeld, and H. Hecht, Temporal-range estimation of multiple objects: Evidence for an early bottleneck. Acta psychologica, 2011. 137(1): p. 76–82.

30. Baurès, R., S.J. Bennett, and J. Causer, Temporal estimation with two moving objects: overt and covert pursuit. Experimental brain research, 2015. 233(1): p. 253–261.

31. Baurès, R., et al., Arrival-time judgments on multiple-lane streets: The failure to ignore irrelevant traffic. Accident Analysis & Prevention, 2014. 65:p. 72–84.

32. Papoulis, A., Probability, random variables, and stochastic processes. 1965.

33. Van Zandt, T., How to fit a response time distribution. Psychonomic bulletin & review, 2000. 7(3): p. 424– 465.

34. Simen, P., et al., Timescale invariance in the pacemaker-accumulator family of timing models. Timing & Time Perception, 2013. 1(2): p. 159–188.

35. Matell, M.S. and W.H. Meck, Neuropsychological mechanisms of interval timing behavior. Bioessays, 2000. 22(1): p. 94–103.

36. Ivry, R.B. and R.M. Spencer, The neural representation of time. Current opinion in neurobiology, 2004. 14(2): p. 225–232.

37. Wearden, J.H., Applying the scalar timing model to human time psychology: Progress and challenges. Time and mind II: Information processing perspectives, 2003:p. 21–39.

38. Gardiner, C.W., Handbook of stochastic methods. Vol. 3. 1985: springer Berlin.

39. Yakimoff, N., N. Bocheva, and L. Mitrani, A linear model for the response time in motion prediction. Acta neurobiologiae experimentalis, 1987. 47(1): p. 55–62.

40. Yakimoff, N., et al., Motion extrapolation performance: A linear model approach. Human factors, 1993. 35(3): p. 501–510.

41. Sharp, R. and H. Whiting, Exposure and occluded duration effects in a ball-catching skill. Journal of Motor Behavior, 1974. 6(3): p. 139–147.

42. Whiting, H. and R. Sharp, Visual occlusion factors in a discrete ball-catching task. Journal of Motor Behavior, 1974. 6(1): p. 11–16.

43. Remington, E. and M. Jazayeri, Late Bayesian inference in sensorimotor behavior. bioRxiv, 2017:p. 130062.

44. Jazayeri, M. and M.N. Shadlen, Temporal context calibrates interval timing. Nature neuroscience, 2010. 13(8): p. 1020.

